# An Integrated Cell Culture - Nanopore Sequencing (ICC-NanoporeSeq) method for the simultaneous detection of multiple enterovirus types and quantification of infectious Enterovirus B

**DOI:** 10.64898/2026.05.15.725335

**Authors:** Aina Astorch-Cardona, Simon Rouault, Tamar Kohn

**Author notes:** Address correspondence to Tamar Kohn.

## Abstract

Enteroviruses (EVs) are ubiquitous contaminants of surface waters, where they can remain infectious for long periods of time. Most methods used for EV monitoring are unable to distinguish between infectious and non-infectious particles or between EV types. Because different types exhibit both distinct environmental persistence and health implications, there is a need for type-resolved infectivity measurements. Here we developed Integrated Cell Culture-Nanopore Sequencing (ICC-NanoporeSeq), a method combining short-term cell culture amplification with Nanopore sequencing of the VP1 gene. The ICC approach was adapted from a previously described ICC-RTqPCR protocol, while the NanoporeSeq workflow was derived from a clinical EV typing protocol and optimized for environmentally circulating EV types. Using samples containing known concentrations of ten EV types, the NanoporeSeq method accurately and reproducibly recovered the original proportions of all EV types after correction of biases. When applied to wastewater extracts, the method detected 45 EV types across five species. Furthermore, type-specific calibration curves generated with ICC-NanoporeSeq enabled quantification of the infectious concentrations of six EV types of the *Enterovirus B* species, allowing a simultaneous and type-resolved assessment of infectivity in mixed samples. Overall, ICC-NanoporeSeq provides a scalable approach for the parallel analysis of multiple EV types. Compared with the predecessor ICC-RTqPCR method, it eliminates the need for multiple type-specific PCR primers and can therefore be readily expanded to include additional EV types.

**IMPORTANCE:** Current methods used to detect EVs in environmental samples generally measure viral genome copies without determining whether viruses remain infectious, limiting their use in public health risk assessment or water quality monitoring. At the same time, available infectivity assays are often labor-intensive and cannot distinguish between different EV types. Here, we developed ICC-NanoporeSeq, a method combining cell culture and Nanopore sequencing to simultaneously quantify the infectious concentrations of multiple EV types in samples containing mixed EV populations. The method provides an efficient and scalable approach for studying EVs in complex environmental matrices. ICC-NanoporeSeq has potential applications in wastewater-based epidemiology, environmental surveillance, and disinfection studies, where understanding the persistence of different EV types simultaneously is crucial.

## INTRODUCTION

Enteroviruses (EVs) are small, non-enveloped RNA viruses that infect mammals. The majority of human-infecting EVs cause mild gastrointestinal or respiratory symptoms^1^, although they can also lead to more severe clinical conditions such as encephalitis, meningitis or myocarditis^2^. Infected individuals shed large quantities of EVs in their feces^3^, leading to the release of viral particles into aquatic environments through spill-overs of raw or partially treated sewage^4^. Due to their high environmental stability, EVs can remain infectious in the aquatic environment for extended periods of time^5,6^, thereby acting as environmental contaminants.

Despite the significant risks posed by EVs and other waterborne viral pathogens, they are still rarely included in routine water quality monitoring programs^7,8^. This is largely due to methodological limitations in detecting infectious viruses. Most existing molecular tools - such as qPCR - quantify viral genomes without distinguishing between infectious and non-infectious particles^5,9^. Yet, only those that remain infectious pose a true sanitary risk. While there exist traditional culture-based infectivity assays that can identify infectious viruses^10^, they require several days to weeks to yield results, and are unable to distinguish between co-circulating EV types^11^ (note that here the term “type” is used in accordance with the current taxonomic nomenclature for EVs, where individual EVs are designated as types rather than serotypes or genotypes^12^). This represents a critical limitation, since different types exhibit variable resistance to environmental and engineered stressors^13–17^, and have distinct public health implications^18^.

To address the need for quantifying type-specific infectious concentrations, Integrated Cell Culture - Reverse Transcription quantitative PCR (ICC-RTqPCR) was developed^11,19^. This approach couples the short-term amplification of viruses in cell culture with type-specific RTqPCR targeting viral RNA. While effective for detecting and quantifying the infectious concentrations of a few selected EV types in a timely manner, this method depends on the prior design and validation of type-specific primers. This limits its scalability and adaptability, particularly when dealing with environmental samples that can contain diverse and dynamic EV populations^20^. Additionally, ICC-RTqPCR workflows are labor-intensive and require frequent optimization for different targets, which challenges large-scale monitoring efforts.

To overcome these limitations, we developed a novel approach that combines cell culture with EV typing using Nanopore sequencing^21^. This method, which we named Integrated Cell Culture - Nanopore Sequencing (ICC-NanoporeSeq), follows the general workflow of ICC-RTqPCR but replaces RTqPCR with high-throughput sequencing of the VP1 gene, the region most commonly used for EV typing^22,23^. Specifically, we adapted an existing Nanopore sequencing typing protocol originally developed for clinical EV strains^21^ to include environmentally circulating EVs. We then evaluated sequencing performance and biases across ten EV types and integrated this approach with cell culture to assess its ability to quantify infectious EVs. By eliminating the need for multiple type-specific RTqPCRs, this method provides a more flexible and informative tool for studying EV dynamics in aquatic systems and for enhancing microbial water quality monitoring within a One Health framework.

## RESULTS

The ICC-NanoporeSeq method developed herein is based on the ICC-RTqPCR method described by Larivé et al^11^, but instead of RTqPCR, it uses Nanopore sequencing for EV typing and quantification. The workflow of ICC-NanoporeSeq (Figure 1A) comprises cell culture amplification of samples containing mixed EV populations, RNA extraction, VP1 amplicon generation, and Nanopore sequencing. The Nanopore sequencing approach (hereafter named NanoporeSeq) was adapted from a protocol by Van Ackeren et al^21^ to type clinical specimens, which we expanded to include environmentally circulating EV strains. Methodological and sequencing data processing details can be found in the Material and Methods section.

**Figure 1.**
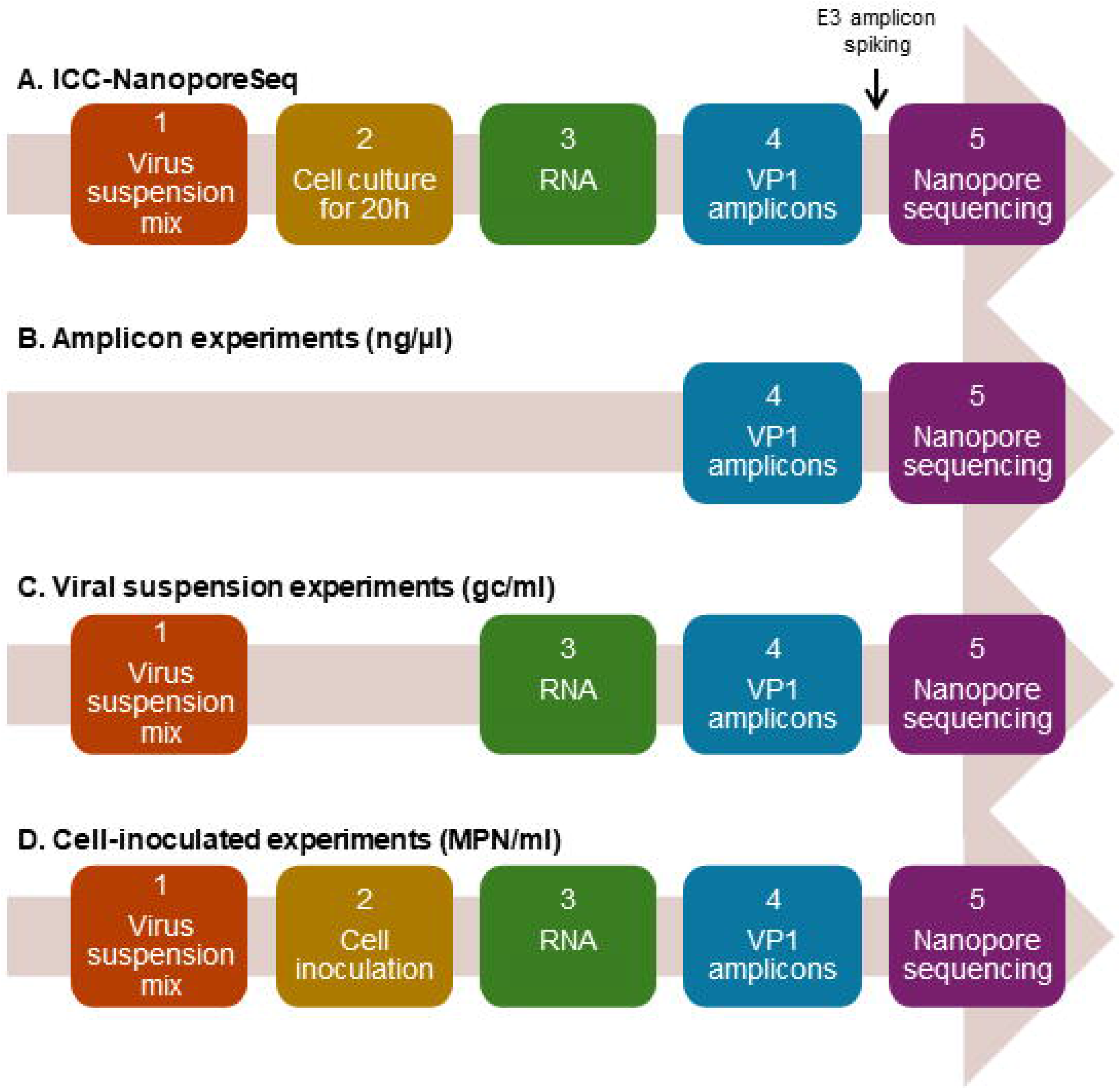
ICC-NanoporeSeq workflow (A) and developmental approach (B-D). The ICC-NanoporeSeq workflow (A) starts by infecting cell cultures with a sample containing mixed EV populations, extracting viral RNA 20 hours post infection, producing VP1 amplicons, adding E3 VP1 amplicon as a spike-in normalizer before library preparation, and sequencing them with Nanopore. The NanoporeSeq method was first tested and optimized on VP1 amplicon mixes (B, amplicon experiments). Such mixes were prepared in terms of ng/µl of cDNA, containing specifically 5.2 ng/µl of VP1 amplicon for each EV type. Subsequently, the method was tested on suspensions containing a mixture of viral particles (C, viral suspension experiments), which were prepared in terms of gc/ml measured by pan-enterovirus RTdPCR. These mixtures contained 108 gc/ml for each EV type. Finally, NanoporeSeq was tested on RD cells inoculated with mixtures of viral particles (D, cell-inoculated experiments). These mixtures were prepared in terms of MPN/ml measured by MPN, specifically containing 102, 103, 104 and 105 MPN/ml for each EV type.

For our experiments, we selected ten EV types from the list proposed by Larivé et al^11^, focusing on those most commonly found in European sewage. Specifically, we included ten EV types belonging to *Enterovirus B* (EV-B): Coxsackieviruses B1, B2, B3, B4, B5, A9 (CVB1, CVB2, CVB3, CVB4, CVB5, CVA9) and Echoviruses 6, 11, 25 and 30 (E6, E11, E25, E30).

### Assessment of NanoporeSeq performance

An initial set of experiments was performed to test whether the adapted NanoporeSeq approach could be used to type and quantify environmentally circulating EV types in suspensions of known virus composition. To this end, we followed a stepwise approach: we first tested and optimized the NanoporeSeq method on VP1 amplicon mixes, then applied it to suspensions containing a mixture of viral particles, and finally to cells inoculated with a mixture of viral particles. In each experiment, we measured relative abundances of all EV types - calculated as the read count for each EV type normalized by the total number of VP1 reads per sample - and compared them to known relative abundances in the original sample. Finally, to test the method’s utility in environmental samples, we applied the NanoporeSeq method to four wastewater samples containing complex and unknown mixtures of EV types.

*Relative abundance analysis.* We first tested NanoporeSeq starting from VP1 amplicon mixes. For these experiments, we extracted RNA and produced VP1 amplicons from stock suspensions of each virus type separately (Figure 1B, amplicon experiments). Subsequently, we composed a mixture containing equal proportions of each amplicon - in terms of ng/µl of complementary DNA (cDNA) - and sequenced the mixture five times. As seen in Figure 2A, the adapted sequencing method successfully captured all EV types considered. The relative abundances of the different types ranged from 4.3% to 22.5% (Figure 2A, D), thus close to the targeted uniform distribution (10%). The sequencing results were highly reproducible across replicates, yielding a mean coefficient of variation (CV) across all replicates and EV types of 3.2%.

**Figure 2.**
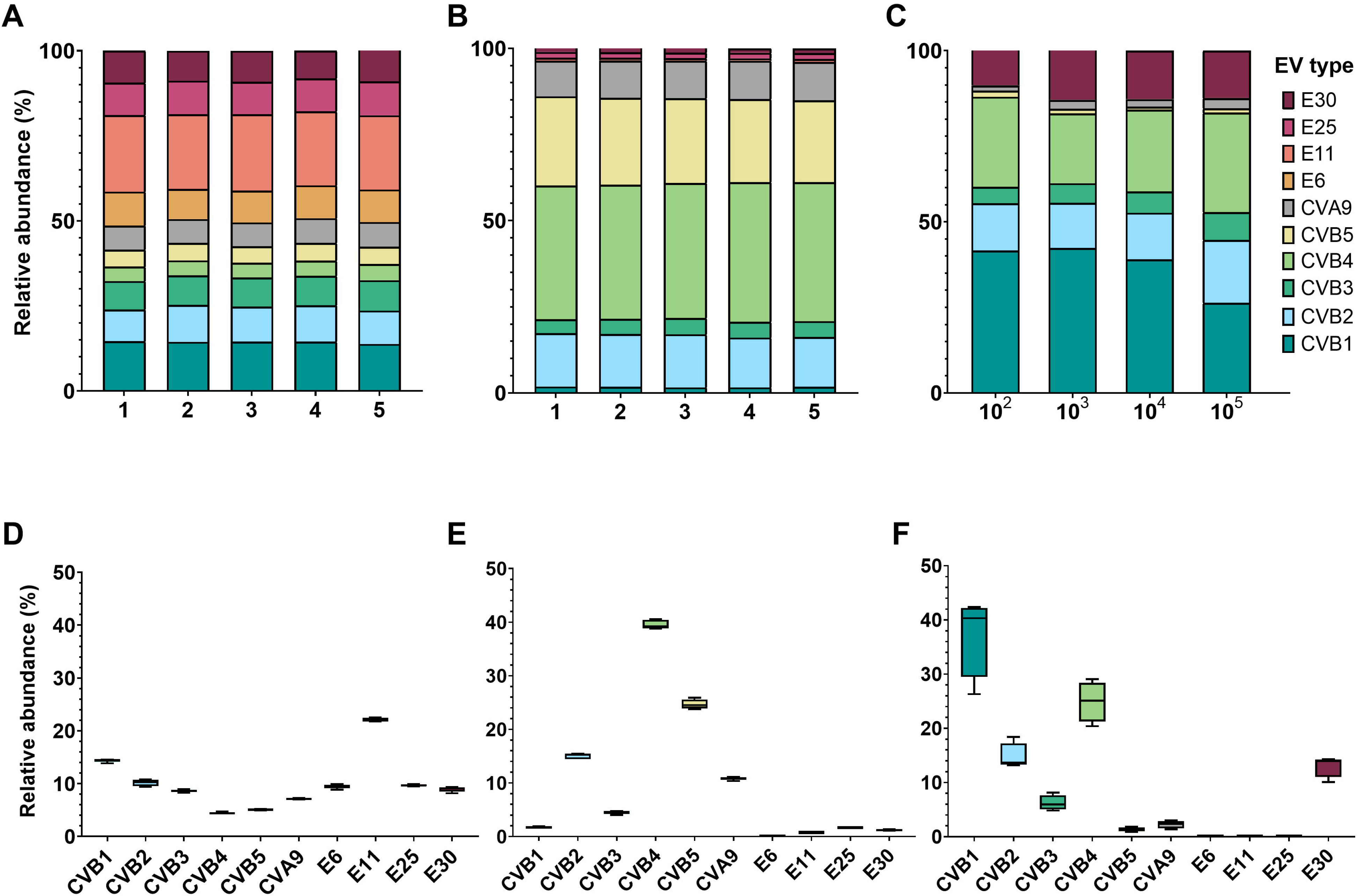
NanoporeSeq protocol performance. A-C: bar plots depicting the relative abundance of each EV type in amplicon (A), viral suspension (B) and cell-inoculated (C) experiments, calculated as the read count of each EV type normalized by the total number of VP1 reads per sample. For amplicon (A) and viral suspension (B) experiments, five replicates were sequenced, while for cell-inoculated (C) experiments, four dilutions were sequenced (MPN/ml). D-F: box plots representing the relative abundance variability of each EV type across replicates for amplicon (D) and viral suspension (E) experiments, or dilutions for cell-inoculated experiments (F). For each EV type, the CV was calculated across all replicates with GraphPad prism (version 11.0.0), and the average of the ten CVs (one per EV type) is reported in the main text.

Next, we prepared a viral suspension containing each EV type in equal proportions - in terms of genome copies (gc)/ml. RNA was extracted from this suspension, and the VP1 amplicon was produced and sequenced five times (Figure 1C, viral suspension experiments). As for amplicon experiments, all EV types could be detected, albeit at widely differing relative abundances, ranging from 0.1% to 40.5% (Figure 2B, E). CVB4 and CVB5 were especially overrepresented in the sequencing results (relative abundance > 23%), whereas E6 and E11 were underrepresented (relative abundance < 1%) (Figure 2B, E). The measured relative abundances were highly reproducible across the five replicate samples, with a mean CV of 5.8%.

Finally, we composed a third viral suspension, this time based on equal infectious titers - in terms of most probable number (MPN)/ml - for each EV type. The suspension was serially diluted three times, resulting in viral titers of 10^2^, 10^3^, 10^4^ and 10^5^ MPN/ml for each EV type. As described in detail in the Material and Methods section, each dilution was inoculated onto a well of confluent Rhabdomyosarcoma (RD) cells. The cells were immediately scraped and subjected to three freeze-thaw cycles, followed by viral RNA extraction, VP1 amplification and sequencing (Figure 1D, cell-inoculated experiments). All EV types remained detectable, though CVB1 and CVB4 were overrepresented (relative abundance > 20%), while E6, E11 and E25 were underrepresented (relative abundance < 1%) (Figure 2C, F). A higher variability was observed compared to the amplicon and viral suspension experiments (mean CV of 38%), which we attribute to the use of serial dilutions of a single virus suspension rather than replicates of equal concentration.

Combined, these results indicate that the adapted NanoporeSeq method could efficiently detect environmental strains of EV, independent of the complexity of the starting suspension. The resulting relative abundances were measured with high reproducibility, though only amplicon experiments accurately reflected the expected ones.

*Bias quantification.* Based on the relative abundances determined above (Figure 2), we quantified three biases associated with the NanoporeSeq protocol: the results of amplicon experiments provide information on bias resulting from library preparation and Nanopore sequencing (sequencing bias). When equal amounts of cDNA from each EV type were combined, we measured relative abundances (Figure 2A, D) that closely matched the expected proportions (10% each type), indicating that this bias is negligible.

When equal amounts of genome copies from each EV type were mixed in viral suspension experiments, the relative abundances obtained after RNA extraction and NanoporeSeq did not match the expected proportions for all EV types (Figure 2B, E). These results therefore revealed a bias associated with the extraction of RNA from the different virus types and their VP1 amplification (here termed extraction/amplification bias). This bias could be quantified as the ratio of raw read counts obtained from viral suspension experiments and amplicon experiments (read ratio; reads_viral suspension_/reads_amplicon_; Supplementary Table S1). Such ratios ranged from 0.01 for E6 to 17.97 for CVB4.

Finally, the results of the cell-inoculated experiments revealed the combined biases arising from the extraction/amplification bias as well as from differences in the genome copies-to-infectivity ratio of different EV types (here termed infectivity bias). The infectivity bias reflects that suspensions used in the cell-inoculated experiments were based on infectious titer, whereas those used in the viral suspension experiments were based on genome copies concentrations. We calculated the type-specific genome copies-to-infectivity ratios based on digital RT-PCR (RTdPCR)-determined genome copies and MPN-derived infectious titers (Supplementary Table S2). The ratios ranged from 0.5 for E25 to 3518 for E30. The < 1 ratio for E25 indicates that the pan-enterovirus RTdPCR assay used herein was not optimal for E25 quantification.

*Bias corrections.* Knowledge of the extraction/amplification and infectivity biases allowed us to reconcile the different relative abundances measured in the amplicon, viral suspension and cell-inoculated experiments (Figure 3). To account for infectivity bias, raw read counts of each EV type obtained from the cell-inoculated experiments were divided by the genome copies-to-infectivity ratios, prior to normalization by the total number of reads. To further correct for the extraction/amplification bias, infectivity bias-corrected reads were then divided by the read ratios prior to normalization. Applying these sequential corrections led to abundance profiles that converged towards those observed in (bias-free) amplicon experiments. Together, these results show that the biases introduced at different stages of the workflow can be sequentially corrected to estimate the initial infectious proportions of the ten EV types present in a sample.

**Figure 3.**
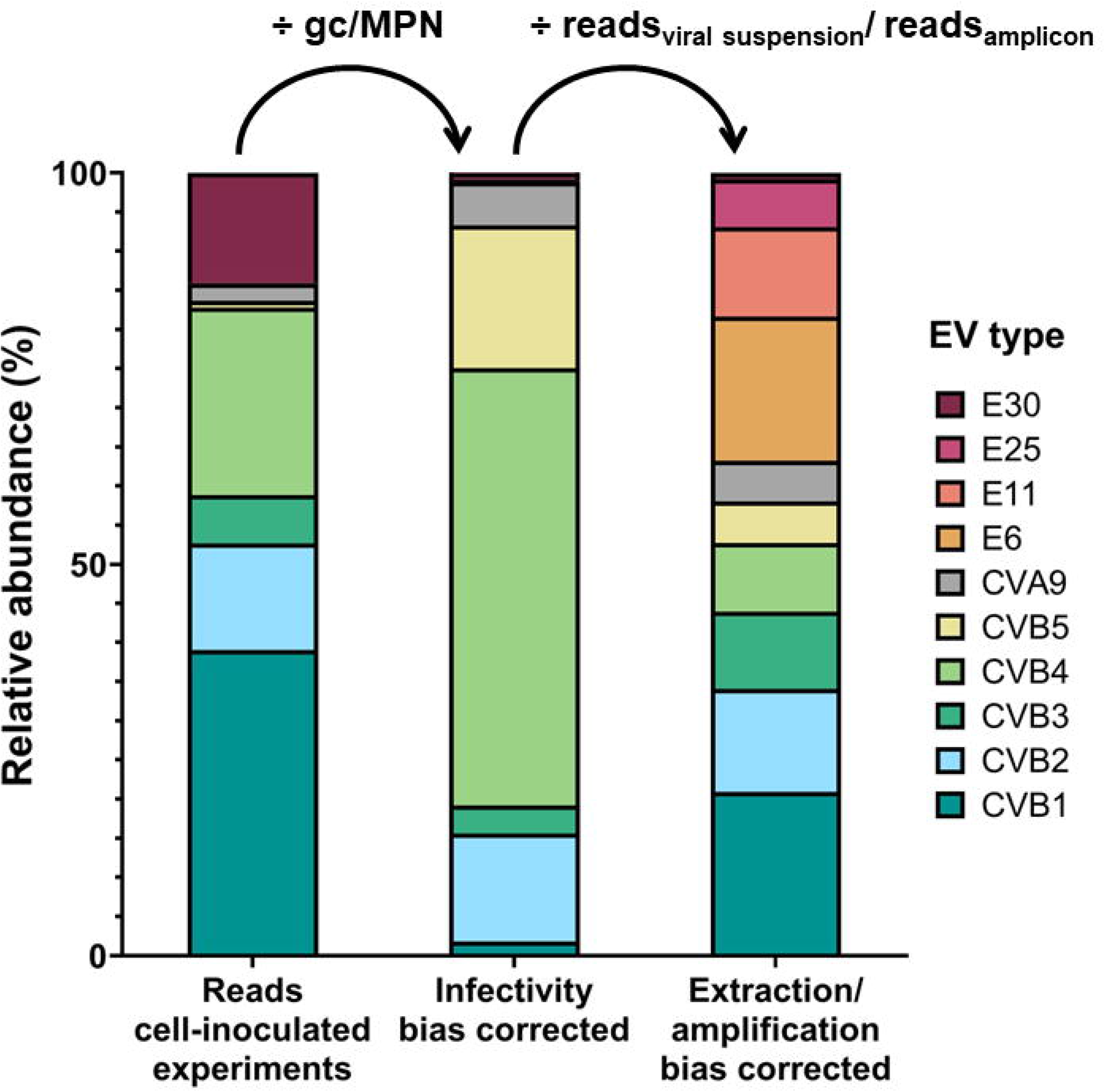
NanoporeSeq method bias corrections. Bar plots depict the relative abundance of reads for each EV type in cell-inoculated experiments (specifically in the dilution containing 104 MPN/ml of each EV type), the relative abundance of reads corrected for the infectivity bias, and for the extraction/amplification bias. To account for the infectivity bias, raw read counts from cell-inoculated experiments were corrected by the genome copies-to-infectivity ratios (Supplementary Table S2). To additionally account for the extraction/amplification bias, the infectivity-corrected reads were subsequently corrected by the read ratio (Supplementary Table S1). Relative abundances were calculated as the read count of each EV type before or after correction, normalized by the total number of VP1 reads.

*Sensitivity analysis*. Two EV types were selected to test the sensitivity of NanoporeSeq: CVB4 and E6. These two types were chosen based on the calculated read ratios (Supplementary Table S1), which revealed that E6 was the most under, and CVB4 the most overrepresented virus when sequencing viral suspensions. Sensitivity tests were first performed in amplicon experiments. A mixture was produced that contained equal proportions of VP1 amplicons from all EV types, except CVB4 or E6, which were added at relative abundances of 1, 0.5 and 0.1%. Measured relative abundances closely matched the expected proportions across the tested range (Figure 4A and C).

**Figure 4.**
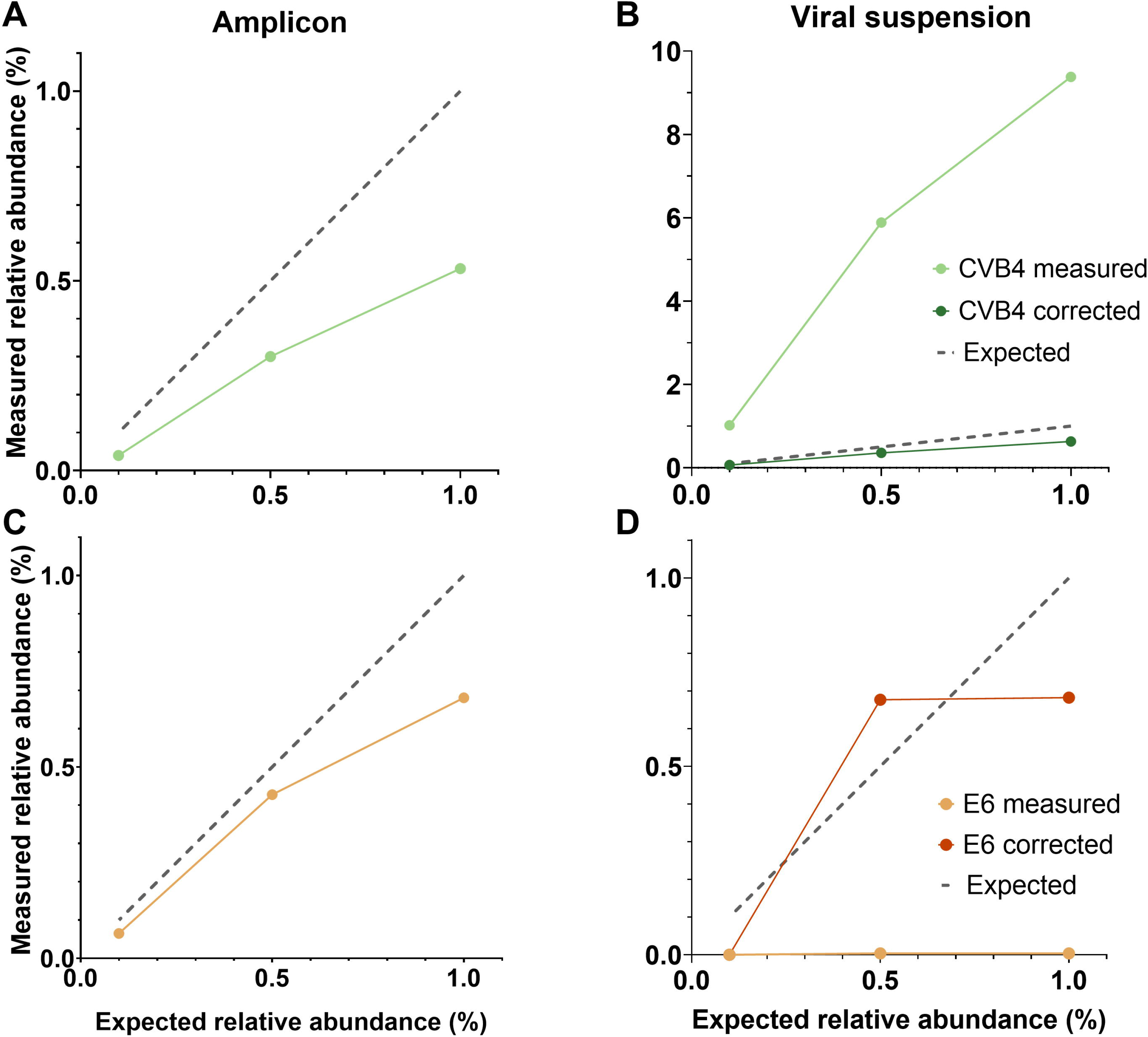
NanoporeSeq method sensitivity analyses for CVB4 and E6. Scatter plot representing the measured vs. the expected relative abundances of CVB4 (A, B) and E6 (C, D) in amplicon (A, C) and viral suspension (B, D) experiments. To account for the extraction/amplification bias in viral suspension experiments, the read ratio (Supplementary Table S1) corrected relative abundance is also shown for both viruses.

Next, sensitivity tests were performed in viral suspension experiments. Resulting relative abundances were strongly biased, with CVB4 being highly overrepresented (Figure 4B) and E6 underrepresented (Figure 4D), as expected from their respective read ratios. These discrepancies were effectively corrected after accounting for extraction/amplification bias, resulting in measured relative abundances closer to the expected values (Figure 4B and D). Together, these results indicate that the NanoporeSeq method can reliably quantify types present at low relative abundances, down to at least 0.1%, even within mixtures dominated by more abundant EV types.

*Application to environmental samples.* To assess whether the NanoporeSeq method could be applied to environmental samples containing unknown mixtures of EV types, we analyzed RNA extracts from four wastewater influent samples collected at different locations across Switzerland (see the Material and Methods section). VP1 amplicons were generated from each RNA extract and sequenced using the NanoporeSeq workflow, as for the viral suspension experiments (Figure 1C).

NanoporeSeq detected not only EV-B, which was the primary focus of the initial method development, but also EV-A, EV-C, and *Rhinovirus A* and *B* (Figure 5). The detection of these additional viral species further demonstrates that the adapted NanoporeSeq workflow is not restricted to EV-B and can be used to characterize a broad range of EVs in complex environmental samples. Quantitative interpretation of relative abundances, however, remains dependent on the availability of type-specific bias corrections. Although correction factors were available for the nine EV types characterized in this study (E25 was not identified in the wastewater samples), additional detected viral types could not be corrected, preventing quantitative comparison of their relative abundances across the viral population.

**Figure 5.**
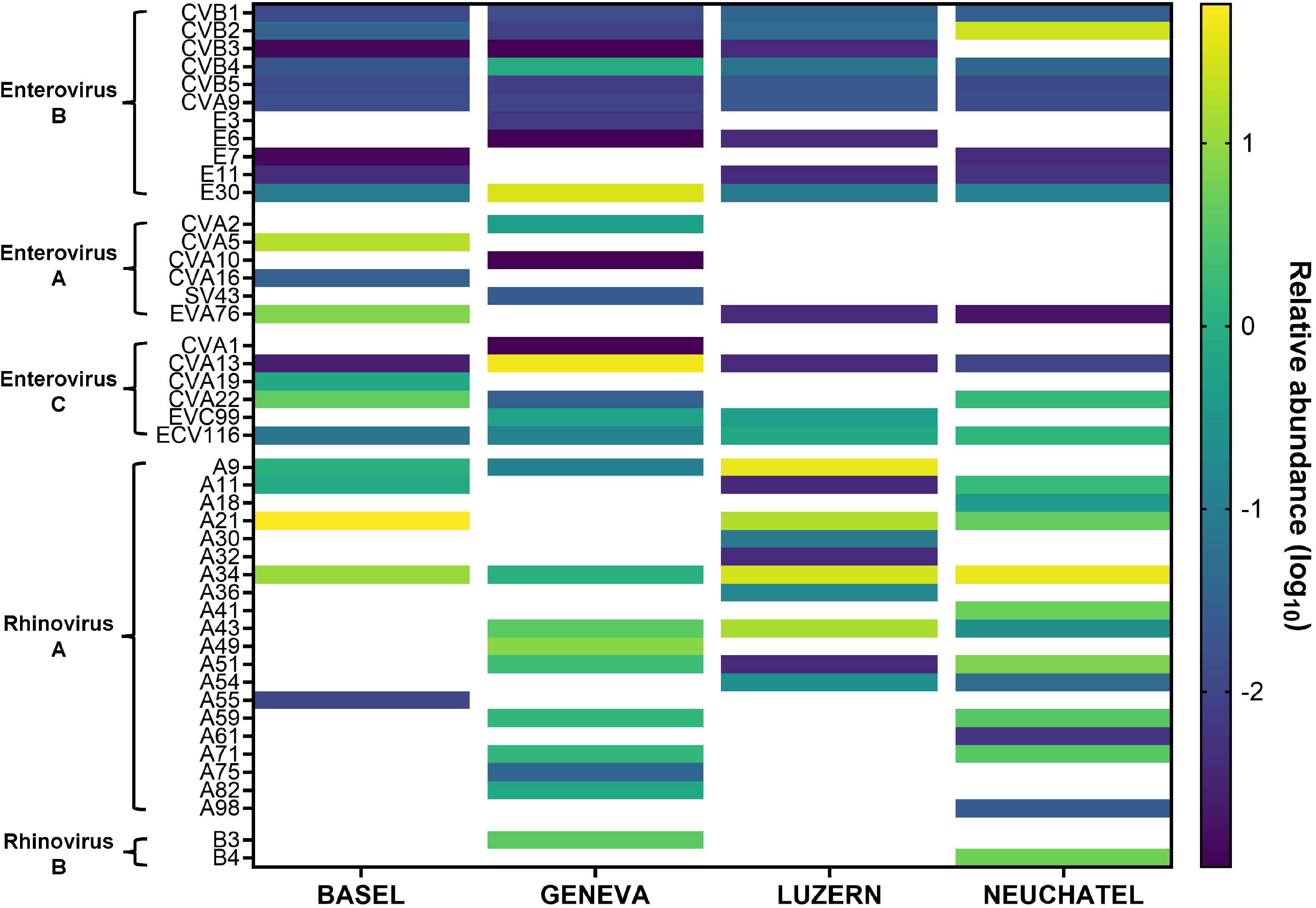
Application of the NanoporeSeq method to wastewater samples. The heatmap shows the log-transformed relative abundances of different EV types detected in four wastewater samples collected in Basel, Geneva, Luzern and Neuchâtel (Switzerland). Relative abundances were calculated as the number of reads assigned to each EV type normalized by the total number of VP1 reads per sample. These values were not corrected for type-specific extraction/amplification biases. Blank cells indicate viral types that were not detected in that sample.

### Calibration of the ICC-NanoporeSeq protocol

Next, we aimed to use the NanoporeSeq protocol in combination with cell culturing, to selectively detect and quantify infectious viruses without the need for prior bias quantification. To do so, we preceded NanoporeSeq with a cell culturing step, to replicate only infectious particles.

To establish EV type-specific calibration curves, we inoculated RD cells with serial dilutions of a virus suspension containing equal infectious titers of all EV types and spanning the dynamic range of the method (see Supplementary Methods and Figure S1). We let the viruses replicate for 20 hours, prior to RNA extraction, VP1 amplification and NanoporeSeq (Figure 1A; ICC-NanoporeSeq). To enable comparison of NanoporeSeq data across samples containing a different starting virus concentration, we amended the samples with a spike-in normalizer (Echovirus 3 (E3) VP1 amplicon) prior to library preparation for sequencing, and normalized all reads against those of E3 (see the Material and Methods section for details).

EV type-specific calibration curves were generated by plotting log-transformed E3-normalized read counts against log-transformed input infectious concentrations (MPN/ml) (Figure 6). Six of the ten EV types replicated in RD cells over the course of 20 hours (see Supplementary Methods and Figure S2). Calibration curves were therefore only generated for CVB5, CVA9, E6, E11, E25, and E30. All six calibration curves showed statistically significant regression slopes (p < 0.05), with R² values ranging from 0.68 to 0.95, except for E30, which showed a lower R² value of 0.37 despite a statistically significant regression (Figure 6, Supplementary Table S3). These calibration curves enable the conversion of sequencing-derived relative abundances after cell culturing into estimates of initial infectious input, thereby providing a quantitative framework for interpreting EV type-specific infectivity for six EV genotypes.

**Figure 6.**
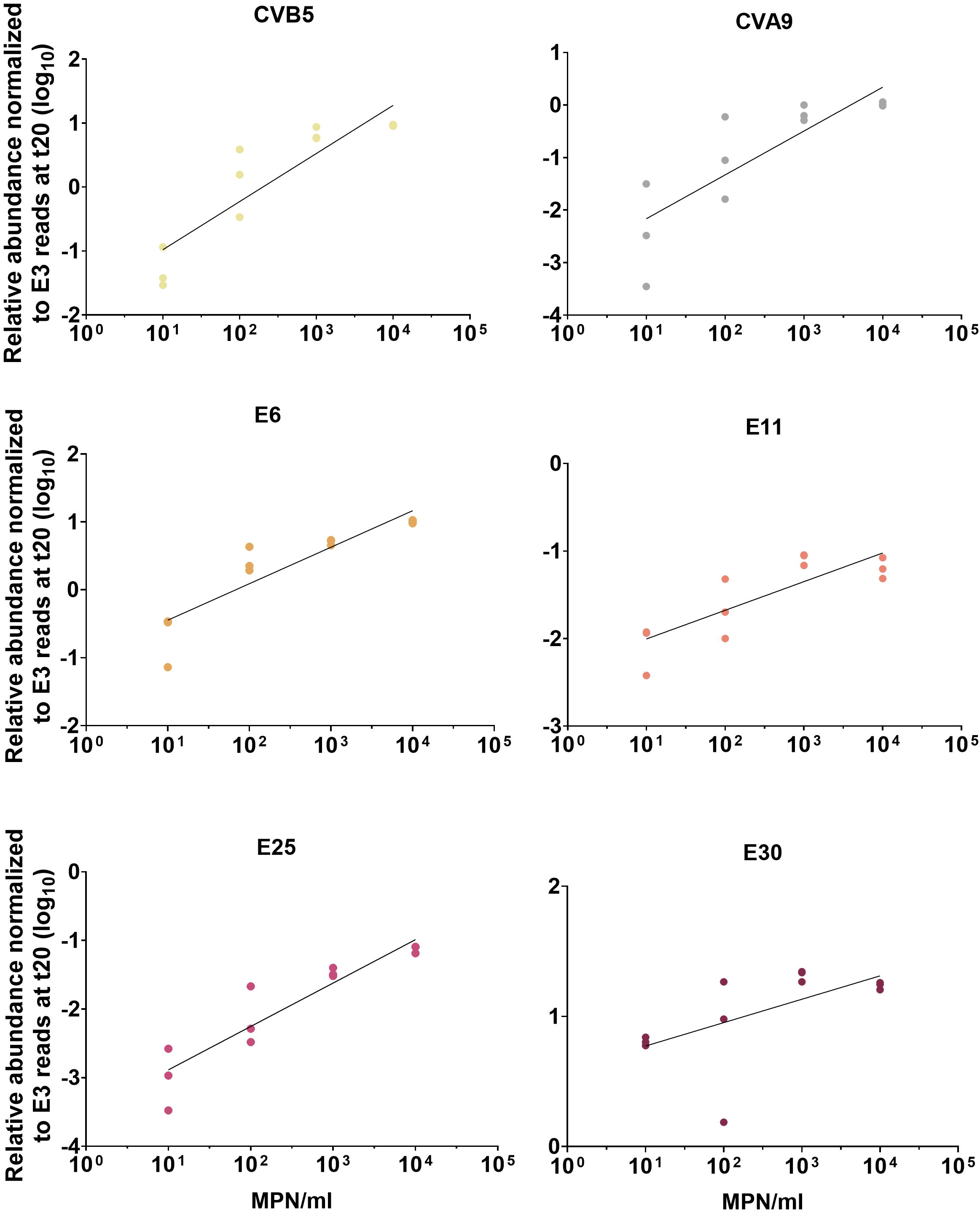
ICC-NanoporeSeq method calibration curves for the six EV types that replicate in RD cells. Calibration curves were obtained by plotting log-transformed, E3-normalized read counts (20 hours post infection, in triplicate) against input log-transformed infectious titers, using four adjacent concentrations. Statistical parameters for calibration curves are given in the Supplementary Material (Supplementary Table S3).

The remaining four EV types (CVB1, CVB2, CVB3, and CVB4) did not show evidence of replication in RD cells (Supplementary Figure S2) and were therefore not included in the calibration analysis. Their inclusion in the ICC-NanoporeSeq approach could potentially be achieved by using alternative cell lines with higher type-specific replication efficiencies (e.g. Buffalo Green Monkey Kidney (BGMK) cells^11^), although this expansion of the method was beyond the scope of this study.

### Comparison of ICC-NanoporeSeq and MPN for infectious concentration quantification

To further evaluate the quantitative performance of ICC-NanoporeSeq, we compared infectious concentrations estimated by ICC-NanoporeSeq with those obtained using the conventional MPN assay. The comparison was performed for the six EV types for which type-specific calibration curves were established (CVB5, CVA9, E6, E11, E25 and E30), using Lake Geneva water as an environmentally relevant matrix. A mixture containing the six EV types at a concentration of 10 □MPN/ml in Lake Geneva water was then analyzed in three independent replicate experiments using the ICC-NanoporeSeq protocol. Infectious concentration estimates were obtained by applying the corresponding type-specific calibration curves (Figure 6) to the E3-normalized read counts obtained after cell culture. Simultaneously, Lake Geneva samples containing 10□MPN/ml of each EV type individually were enumerated by MPN in three replicate experiments.

After accounting for the ten-fold difference in starting concentrations between the two approaches, ICC-NanoporeSeq estimates were overall comparable to the concentrations measured by MPN (Figure 7). ICC-NanoporeSeq consistently estimated infectious concentrations close to the expected input for all six EV types, whereas MPN measurements showed greater variation. The estimates obtained by the two approaches were relatively similar for CVB5, CVA9 and E11, while E6, E25 and E30 showed greater differences between the two methods, although these remained below one order of magnitude. Taken together, these results demonstrate that ICC-NanoporeSeq provides infectious concentration estimates that are consistent with the expected concentrations and comparable to those obtained by MPN.

**Figure 7.**
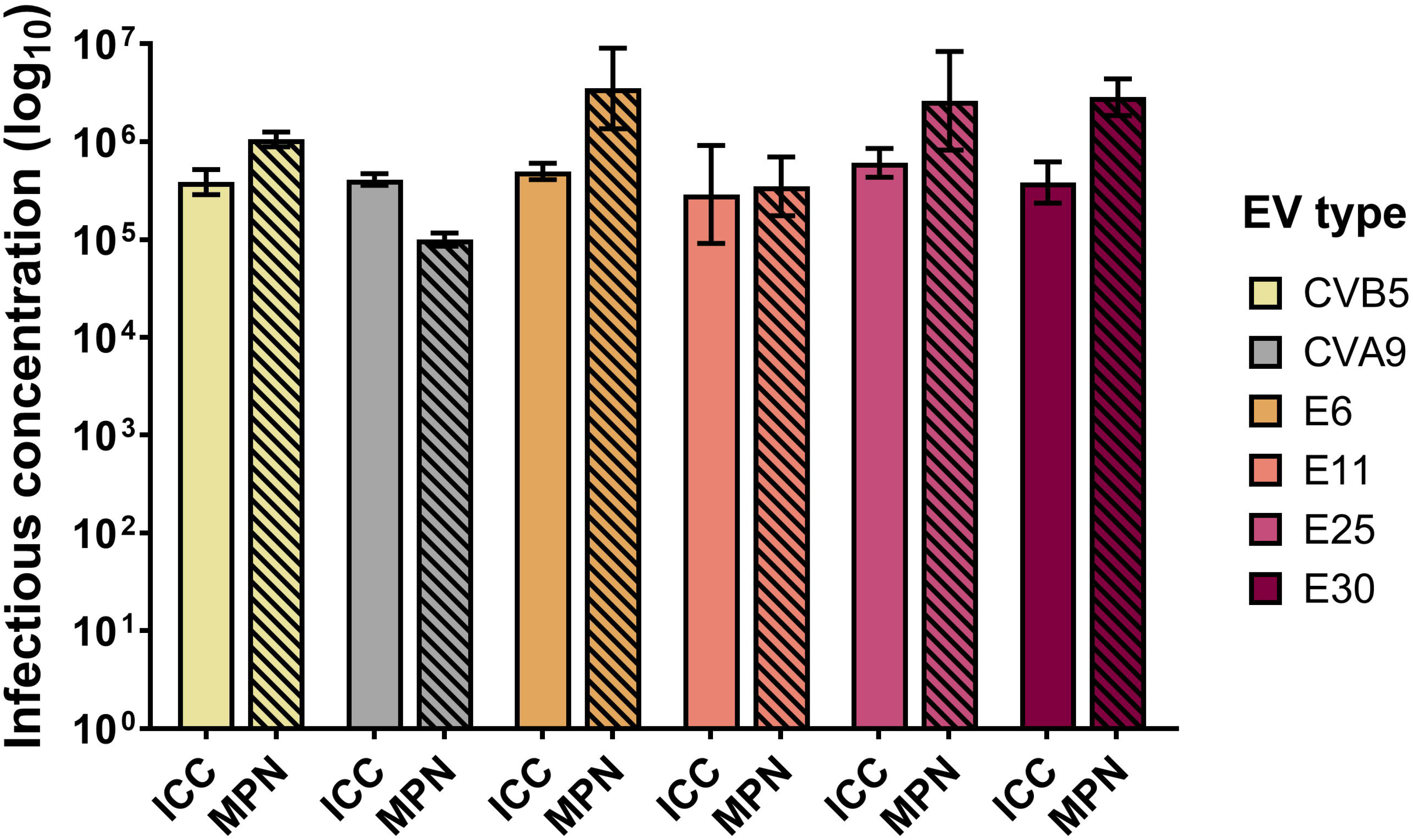
Log-transformed infectious concentrations of CVB5, CVA9, E6, E11, E25 and E30 estimated using ICC-NanoporeSeq (ICC) and MPN in Lake Geneva water. For ICC-NanoporeSeq, the six EV types were simultaneously spiked at a concentration of 10 □ MPN/ml per type, and infectious concentrations were estimated using the corresponding type-specific calibration curves. For MPN, each EV type was spiked individually at 10 □ MPN/ml. For visualization and comparison with MPN, ICC-NanoporeSeq estimates were multiplied by 10 to account for the ten-fold difference in starting concentrations between the two approaches. Bars represent the mean infectious concentration of three independent replicates (with duplicate samples for ICC-NanoporeSeq), and error bars indicate standard deviation.

## DISCUSSION

The results obtained during the development and optimization of the ICC-NanoporeSeq method demonstrate that NanoporeSeq can be used as a robust quantitative tool to resolve the composition of samples containing mixed populations of EVs when appropriate bias corrections are applied (Figure 3). Correction for extraction/amplification bias allows for the quantification of total (infectious plus inactivated) EV type distribution. In experiments using virus types with well-characterized genome copies-to-infectivity ratios, e.g., disinfection experiments conducted in laboratory settings, correction for infectivity bias can furthermore resolve the composition of infectious EV types. In environmental samples, where the viral population is diverse and type-specific biases may not have been fully established, NanoporeSeq can still be used to identify the EV types present and characterize their diversity.

The integration of a cell culture amplification step enables the assessment of infectivity in samples of unknown infectivity bias. In line with the principles of ICC-RTqPCR approaches^11,19,24^, the increase in viral genome copies following cell infection is proportional to the initial concentration of infectious virus, which allows to correlate the measured signal to the infectious input within a defined dynamic range. In the present study, we first assessed the replication of the ten EV types in RD cells over a 20-hour cell culture amplification period (Supplementary Figure S2), and only EV types showing evidence of replication were included in the calibration experiments. Calibration curves using RD cells therefore enable the conversion of ICC-NanoporeSeq-derived reads into estimates of infectious input for a subset of six EV types (Figure 6).

Four EV types did not show sufficient replication in RD cells over the 20-hour cell culture amplification period to be included in the calibration curves (Supplementary Figure S2). As previously observed in ICC-based methods, this limitation likely reflects differences in replication efficiency and host cell compatibility. While Larivé et al^11^ suggested that calibration may need to be established for each experiment, the present results indicate that calibration robustness is also driven by EV type-host interactions rather than experimental variability alone. Expanding the method to include additional cell lines such as BGMK or HeLa could improve its coverage across a broader range of EV types within and beyond the EV-B species. However, some EV types may not readily replicate in commonly used cell cultures, and the use of multiple cell lines would increase experimental complexity.

Beyond method validation, the ICC-NanoporeSeq approach provides a versatile framework for EV research and surveillance. In the absence of the cell culture amplification step, the method can be applied to characterize EV type composition in environmental samples, supporting applications such as wastewater-based epidemiology and environmental monitoring. Our analysis of four wastewater samples demonstrates the ability of NanoporeSeq to detect and characterize diverse EVs, including members of the EV-A, EV-B, EV-C and *Rhinovirus A* and *B*. As environmental EV populations are diverse and dynamic, continued expansion and updating of the reference database with sequences from environmentally circulating strains will be important to ensure an accurate characterization of EV diversity.

When combined with the infectivity assay, ICC-NanoporeSeq enables a simultaneous assessment of infectivity of six different types, making it particularly suitable for disinfection and inactivation studies, notably in research related to viral environmental persistence. In our comparison with MPN, six EV types were quantified individually by MPN and simultaneously as a mixture by ICC-NanoporeSeq, with comparable infectious concentrations obtained using the two approaches (Figure 7). The ability to analyze multiple types in parallel represents a substantial reduction in time and laboratory resources compared to conventional single-type assays, which would otherwise require multiple independent experiments.

Despite these advantages, several limitations should be considered. Biases introduced during RNA extraction and VP1 amplicon production require careful correction, as demonstrated in this study. In addition, not all EV types replicate efficiently in RD cells, limiting quantitative analysis for some viruses and highlighting the importance of host cell selection in ICC-based approaches. Ensuring a broader coverage of EV types may require the use of additional cell lines, which would increase experimental complexity and represent a trade-off with the scalability of the approach. From a practical perspective, ICC-NanoporeSeq also requires Nanopore sequencing infrastructure and bioinformatic expertise in addition to the cell culture and molecular biology procedures already required for MPN or ICC-RTqPCR. Addressing these limitations, as well as further standardizing both wet-lab and bioinformatic workflows^25^, will be essential to improve the robustness and general applicability of the method.

## MATERIAL AND METHODS

### Viral stock suspensions

One environmental isolate was obtained for nine of the ten selected EV types. Coxsackieviruses B4 (CVB4, GenBank accession number MG845888) and B5 (CVB5, GenBank accession number MG845891) were obtained from Minneapolis and Lausanne sewage, respectively, using previously described methods^13^. The remaining types were kindly provided by Soile Blomqvist and Carita Savolainen-Kopra from the Finnish National Institute for Health and Welfare (Helsinki, Finland), and included Coxsackieviruses B1, B2, B3, A9 (CVB1, CVB2, CVB3, CVA9) and Echoviruses 6, 25 and 30 (E6, E25, E30), as well as Echovirus 3 (E3), which was used as a spike-in control in the ICC-NanoporeSeq protocol. For Echovirus 11 (E11), the ATCC-Gregory lab strain (GenBank accession number X80059) was used.

Viral stocks of each genotype were prepared as detailed by Larivé et al^11^. Briefly, all Coxsackieviruses B were propagated on BGMK cells (provided by the Spiez Laboratory, Spiez, Switzerland), while all Echoviruses - and CVA9 - were propagated on RD cells (ATCC CCL-136, Virginia, USA). Both cell lines were grown at 37°C in a 5% CO_2_ atmosphere, using Minimum Essential Medium (MEM, Life Technologies, CA, USA) for BGMK cells, and Dulbecco’s Modified Eagle Medium (DMEM, Life Technologies, CA, USA) for RD cells. Both media were supplemented with 10% Fetal Bovine Serum (FBS, Life Technologies, CA, USA) and 1% penicillin/streptomycin (Life Technologies, CA, USA). For cell maintenance, the media was supplemented with 2% FBS instead of 10%.

Confluent flasks of BGMK or RD cells were infected with the virus isolate in maintenance media and incubated at 37°C until full cytopathic effect (CPE) was observed. The cell lysate was then subjected to three freeze-thaw cycles, followed by centrifugation at 3500 *g* for 10 minutes. The resulting supernatant was subjected to buffer exchange to replace the culture media with phosphate-buffered saline (PBS) using 100 kDa Amicon Ultra Centrifugal Filters (Merck Millipore, MA, USA). Finally, the viral stock was aliquoted and stored at −80°C. Viral stocks were titrated on their respective cell lines via endpoint dilution in 96-well plates, and their infectious concentrations were determined using the MPN method^26^ with 5 replicates per dilution. Viral genome copy numbers were determined in each stock by RTdPCR for pan-enteroviruses, as described below (see *RNA extractions and pan-enterovirus RTdPCR*). All experiments in this study were performed with the same generation of viruses except for E30, for which two different generations were used.

### RNA extractions and pan-enterovirus RTdPCR

All viral RNA extractions were performed using the QIAamp Viral RNA Mini Kit (Qiagen, Hilden, Germany), following the manufacturer’s instructions. Viral RNA was extracted from 140 µl of sample, eluted twice in 40 µl of elution buffer in order to increase RNA yields, and stored at −20°C prior to VP1 amplicon production or RTdPCR.

Pan-enterovirus RTdPCR was performed using the QIAcuity Digital PCR System (QIAcuity One 2plex Device), the QIAcuity Nanoplates 8.5k 24- or 96-well and the QIAcuity OneStep Advanced Probe Kit (all from Qiagen, Hilden, Germany) as previously described^27^. In brief, each RTdPCR reaction mixture contained 3 μl of QIAcuity OneStep Advanced Probe Kit Mastermix 4x, 0.12 μl of RT enzyme, 1.5 μl of GC enhancer, 0.6 μl of primers and 0.3 μl of probe at a working concentration of 20 μM, 1.88 μl of DNase-free water and 4 μl of template RNA in a final volume of 12 μl. A DNase-free water negative control was included in each RTdPCR run. The forward and reverse primers for pan-enteroviruses were 5’-CCTCCGGCCCCTGAATG-3’ and 5’-ACCGGATGGCCAATCCAA-3’, respectively. The probe used was 5’-HEX-CGGAACCGACTACTTTGGGTGTCCGT-BHQ1-3’ (Microsynth AG, Balch, Switzerland). The RTdPCR program consisted of a 40-minute RT step at 50°C and a 2-minute RT enzyme inactivation step at 95°C, followed by 45 cycles of a 15-second denaturation step at 95°C and a 1-minute annealing and extension step at 60°C. QIAcuity Software Suite (version 3.2.0.0) (Qiagen, Hilden, Germany) was used for experimental parameter set-up and data analysis.

### Integrated cell culture (ICC)

The ICC protocol was adapted from the ICC-RTqPCR method described by Larivé et al^11^. RD cells were seeded in 6-well plates and allowed to reach full confluency. After removing the growth media, duplicate wells were inoculated with 1 ml of viruses in maintenance media and were incubated for 20 hours at 37°C before being scraped and frozen at −20°C. After three freeze-thaw cycles, the recovered viruses and cells were centrifuged for 10 minutes at 1000 *g* to remove cell debris, the supernatant was transferred into a clean tube and RNA was extracted from 140 μL of supernatant as described above (see *RNA extractions and pan-enterovirus RTdPCR*). Finally, VP1 amplicons were produced and sequenced following the protocols described below (see *VP1 amplicon production and NanoporeSeq*).

### VP1 amplicon production and NanoporeSeq

EV typing was performed via VP1 amplicon sequencing, following a protocol adapted from Van Ackeren et al^21^. To produce the VP1 amplicon, a region within the VP1 gene was amplified as previously detailed by Nix et al^23^ and the WHO enterovirus surveillance guide^28^, by performing a reverse transcription step followed by a semi-nested PCR. All primers were synthesized at Microsynth AG (Balch, Switzerland).

Briefly, a mixture containing 0.5 mM dNTPs and 0.1 µM of each of four RT primers (AN32, AN33, AN34, and AN35; Supplementary Table S4) was prepared. An 8 µl aliquot of this mixture was added to 5 µl of RNA template in a final reaction volume of 13 µl and was incubated at 65°C for 5 minutes to denature secondary RNA structures. Complementary DNA (cDNA) was then generated using the SuperScript IV Reverse Transcriptase (Thermo Fisher Scientific, MA, USA) according to the manufacturer’s protocol.

Subsequently, a first PCR was performed to produce an amplicon of approximately 800 bp. This reaction was carried out in a total volume of 25 µl and contained 0.05 U/µl AmpliTaq Polymerase, 1x PCR Buffer I (Thermo Fisher Scientific, MA, USA), 0.2 mM dNTPs, 1 µM of each forward and reverse primer (SO224 and SO222, respectively; Supplementary Table S4), and 5 µl of cDNA. The cycling conditions were as follows: initial denaturation at 95°C for 2 minutes, followed by 35 cycles of 95°C for 15 seconds, 42°C for 30 seconds, and 72°C for 45 seconds, with a final extension at 72°C for 5 minutes.

A second, semi-nested PCR was then performed to produce an amplicon of approximately 400 bp. This reaction was performed in a final volume of 50 µl containing 0.05 U/µl AmpliTaq Polymerase, 1x PCR Buffer I, 0.8 µM of each forward and reverse primer (AN89 and AN88, respectively; Supplementary Table S4), and 1 µl of the product from the first PCR. The cycling conditions were: initial denaturation at 95°C for 2 minutes, followed by 38 cycles of 95°C for 15 seconds, 60°C for 30 seconds, and 72°C for 45 seconds, with a final extension at 72°C for 5 minutes.

The final amplicon was purified using 200% volume of Agencourt AMPure XP beads (Beckman Coulter, CA, USA), washed twice with 70% ethanol, and eluted in 30 µl of 10 mM Tris-HCl pH 8.0 with 50 mM NaCl. The cDNA concentration of the purified amplicon was quantified using the Qubit 1x Broad Range dsDNA Assay on a Qubit 4 fluorometer (Thermo Fisher Scientific, MA, USA). VP1 amplicons of each viral stock were Sanger sequenced (Microsynth AG, Balch, Switzerland) and typed using the online EV genotyping tool^29^. Sequences for each EV type can be found in the Supplementary Material.

Nanopore sequencing libraries were prepared using the Native Barcoding Kit 24 V14 (SQK-NBD114.24, ONT, Oxford, UK), following the manufacturer’s instructions. Briefly, VP1 amplicons were end-prepped, barcoded and ligated to sequencing adapters. For each run, up to 24 samples were pooled together and sequencing libraries were prepared using at least 100 fmol (corresponding to 25 ng) of DNA. Each library was loaded on a R10.4.1 flow cell (FLO-MIN114, ONT, Oxford, UK) and was sequenced for 24 hours on a MinION Mk1C device (ONT, Oxford, UK).

Raw Pod5 reads were basecalled using the Oxford Nanopore high accuracy basecaller Dorado (v0.9.1, https://github.com/nanoporetech/dorado). Basecalled reads were processed using a custom pipeline based on the publicly available Snakemake script developed by Van Ackaren et al^21^ to type EVs from clinical samples (https://github.com/medvir/ONT_amplicon/releases/tag/v1.0, last accessed on 20.02.2026). The original workflow concatenated reads from multiple FASTQ files per sample, trimmed barcode and primer sequences to extract the amplicons, mapped them to a homemade reference database of 538 EV sequences using minimap2 and created a consensus sequence for each sample. Amplicon extraction was the only read/quality filtering step and allowed no mismatches while supporting degenerate bases. Because this step was stringent, all reads passing the filter were retained for downstream analysis. For the purposes of this study, we implemented the following modifications: (i) the original reference database was replaced with a custom database comprising the original 538 EV sequences and 213 more sequences from environmental EV isolates, obtained both from NCBI and from Sanger sequencing of the viral stocks used; and (ii) the consensus sequence generation step was removed to focus exclusively on quantifying the number of reads per type. All code and modified scripts are available at 10.5281/zenodo.20154852.

### NanoporeSeq application to environmental samples

The NanoporeSeq method was applied to RNA extracts from four wastewater samples kindly provided by Tim Julian and Nadja Widrig (Swiss Federal Institute of Aquatic Science and Technology - EAWAG, Switzerland). Influent wastewater samples were collected at wastewater treatment plants in Basel, Geneva, Luzern and Neuchâtel (Switzerland). Sample collection, processing and RNA extraction were performed as previously described^30,31^. RNA extracts were subsequently used for VP1 amplification and Nanopore sequencing following the workflow described in the main text (see *VP1 amplicon production and NanoporeSeq*). Viral types were identified based on the resulting VP1 sequences, and their relative abundances were determined as the number of reads assigned to that type normalized by the total number of VP1 reads per sample.

### ICC-NanoporeSeq calibration

To determine if the ICC-NanoporeSeq method was able to yield quantitative information on infectious EV concentrations, we first tested all EV types used in this study for their ability to replicate in RD cells (see Supplementary Methods for details). Of the ten types included, six were found to replicate (CVB5, CVA9, E6, E11, E25, and E30). For these six virus types we then established type-specific calibration curves. Specifically, we prepared four calibrant suspensions in maintenance media containing 10^1^, 10^2^, 10^3^ and 10^4^ MPN/ml of each virus type. These infectious concentrations fell within the dynamic range of the ICC-RTqPCR method published by Larivé et al^11^ and was confirmed for one virus type (E11) herein as described in the Supplementary Material.

The calibrant samples (in duplicate) were then inoculated in RD cells and subjected to the ICC-NanoporeSeq protocol described above. While calibrant suspensions had different virus concentrations, they all contained the same proportion of the different EV types. Therefore, after amplification of the VP1 gene, 10 µl of each sample were amended with 1 µl of E3 VP1 amplicon. Hereby, the constant concentration of the E3 VP1 amplicon across samples served as a spike-in normalizer that would allow to assess differences in the amount of cDNA obtained from different starting virus concentrations in the calibrant suspensions. Finally, the amount of cDNA in the spiked suspensions was measured by Qubit and 52 ng were used for subsequent Nanopore library preparation. NanoporeSeq read counts obtained for each EV type were normalized to the corresponding E3 read counts, allowing comparison of relative abundances across samples.

Calibration curves for each EV type were obtained by analyzing the averaged log-transformed, E3-normalized read counts against the corresponding log-transformed infectious input concentration of the calibrants and fitting a linear regression. Statistical parameters for each calibration curve were determined using GraphPad Prism (version 11.0.0). Each calibration curve included four calibrant concentrations in triplicate.

### ICC-NanoporeSeq and MPN comparison in Lake Geneva water

One litre of Lake Geneva water was collected in a sterile glass bottle at the shore of Pelican Beach (Saint-Sulpice, Switzerland), using a sampling pole. Once in the laboratory, the sample was filtered through 0.8 µm mixed cellulose ester filters (Merck Millipore, MA, USA) and 15 ml aliquots were transferred into sterile 50 ml centrifuge tubes. For MPN analysis, each of the six EV types included in the calibration experiments (CVB5, CVA9, E6, E11, E25 and E30) was spiked individually into three independent 15 ml aliquots to obtain a final concentration of 10 □ MPN/ml. For ICC-NanoporeSeq analysis, a mixture containing the ten EV types at a final concentration of 10 □ MPN/ml for each type was prepared and added to three independent 15 ml aliquots.

Immediately after spiking, 1 ml samples were collected from each 15 ml aliquot and frozen at −20°C until analysis. Samples were subsequently analyzed by MPN or ICC-NanoporeSeq as described in the corresponding sections of the Materials and Methods (see *Viral stock suspensions* for MPN and *ICC-NanoporeSeq calibration* for ICC-NanoporeSeq). ICC-NanoporeSeq infectious concentration estimates were obtained for the six EV types by applying the corresponding type-specific calibration curves to the E3-normalized read counts.

## Data availability

The datasets generated during this study have been deposited in Zenodo and are publicly available at 10.5281/zenodo.22749196.

## ACKNOWLEDGMENTS

This work was supported by the Swiss National Science Foundation Water4All project no. 20WA21_216704. The funders had no role in study design, data collection and interpretation, or the decision to submit the work for publication. We thank Tim Julian and Nadja Widrig for providing wastewater extracts, Sujin Shin for assistance with data analysis and interpretation and Laurine Mancardi and Loïs Bendera for technical assistance.

## Notes

https://doi.org/10.5281/zenodo.22749196

### Competing Interest Statement

The authors have declared no competing interest.

### Summary of Updates

Most importantly, we have: changed the title of the manuscript to better reflect that infectious virus quantification was only done for types of the Enterovirus B species (whereas detection is possible for multiple species); conducted analyses in wastewater samples to demonstrate the applicability of the sequencing method to real samples. Specifically, we demonstrate that the method can detect many different enterovirus types from several species in real samples; validated the performance of the ICC-NanoporeSeq against traditional culturing methods for the quantification of six infectious virus types that replicate in RD cells; expanded our calibration curves for greater robustness. Because the manuscript includes new experimental data, we have added an additional author (Simon Rouaoult) who conducted the necessary analyses.

