## Supplementary Material for "An Integrated Cell Culture - Nanopore Sequencing (ICC-NanoporeSeq) method for the simultaneous detection of multiple enterovirus types and quantification of infectious Enterovirus B"

**Supplementary Table S1.** Read ratios for each EV type and data used for their calculation. Read ratios were calculated as the average of raw reads (not normalized) obtained in the five replicate samples sequenced in amplicon experiments, divided by the average of raw reads obtained in viral suspension experiments.

| **EV type** | **Amplicon experiments**  **(average raw reads)** | **Viral suspension experiments**  **(average raw reads)** | **Read ratio**  **(reads_viral suspension_/reads_amplicon)_** |
| --- | --- | --- | --- |
| CVB1 | 6789.6 | 1651.0 | 0.24 |
| CVB2 | 4809.8 | 14158.8 | 2.97 |
| CVB3 | 4084.6 | 4194.0 | 1.03 |
| CVB4 | 2084.8 | 37248.6 | 17.97 |
| CVB5 | 2404.0 | 23339.8 | 9.73 |
| CVA9 | 3354.4 | 10188.0 | 3.04 |
| E6 | 4468.0 | 57.8 | 0.01 |
| E11 | 10428.0 | 701.6 | 0.07 |
| E25 | 4577.2 | 1553.2 | 0.34 |
| E30 | 4212.8 | 1148.0 | 0.28 |

**Supplementary Table S2.** Genome copies-to-infectivity ratios for each EV type and data used for their calculation. Genome copies-to-infectivity ratios were calculated as the genome copies concentration of each EV type determined by pan-enterovirus RTdPCR, divided by the infectious titer of each EV type determined by MPN.

| **EV type** | **Genome copies concentration**  **(gc/ml)** | **Infectious concentration (MPN/ml)** | **Genome copies-to-infectivity ratio (gc/MPN)** |
| --- | --- | --- | --- |
| CVB1 | 5.06 x 10^9^ | 1.19 x 10^7^ | 424.83 |
| CVB2 | 2.31 x 10^10^ | 1.19 x 10^9^ | 19.40 |
| CVB3 | 1.57 x 10^10^ | 4.64 x 10^8^ | 33.76 |
| CVB4 | 7.12 x 10^9^ | 8.50 x 10^8^ | 8.38 |
| CVB5 | 3.16 x 10^10^ | 3.34 x 10^10^ | 0.95 |
| CVA9 | 7.89 x 10^9^ | 1.03 x 10^9^ | 7.66 |
| E6 | 2.72 x 10^10^ | 5.47 x 10^9^ | 4.97 |
| E11 | 1.63 x 10^10^ | 1.59 x 10^10^ | 1.03 |
| E25 | 3.66 x 10^8^ | 7.03 x 10^8^ | 0.52 |
| E30 | 1.63 x 10^10^ | 4.64 x 10^6^ | 3518.35 |

**Supplementary Table S3.** ICC-NanoporeSeq calibration curve statistics. Simple linear regression statistics (slope, intercept, p-value and R^2^) were performed on GraphPad Prism (version 11.0.0). Significant p-values are marked with a *. Calibration curves are only shown for those EV types that efficiently replicate in RD cells over the course of 20 hours (Supplementary Figure S2 and Supplementary Table S5).

| **EV type** | **Slope** | **Intercept** | **p-value** | **R^2^** |
| --- | --- | --- | --- | --- |
| CVB5 | 0.75 | -1.73 | <0.0001* | 0.81 |
| CVA9 | 0.83 | -3.00 | 0.0006* | 0.70 |
| E6 | 0.54 | -0.98 | <0.0001* | 0.81 |
| E11 | 0.33 | -2.33 | 0.001* | 0.68 |
| E25 | 0.63 | -3.52 | <0.0001* | 0.86 |
| E30 | 0.18 | 0.59 | 0.03* | 0.37 |

**Supplementary Table S4.** Primers used for VP1 amplicon production, from the WHO enterovirus surveillance guide^1^. Standard International Union of Biochemistry nucleotide ambiguity codes are used: I = Deoxyinosine, N = G, A, T or C, Y = C or T, W = A or T, R = A or G.

| **Primer name** | **Primer function** | **Primer sequence** |
| --- | --- | --- |
| AN32 | RT | 5’ GTY TGC CA 3’ |
| AN33 | RT | 5’ GAY TGC CA 3’ |
| AN34 | RT | 5’ CCR TCR TA 3’ |
| AN35 | RT | 5’ RCT YTG CCA 3’ |
| SO224 (F) | First PCR | 5’ GCI ATG YTI GGI ACI CAY RT 3’ |
| SO222 (R) | First PCR | 5’ C ICC IGG IGG IAY RWA CAT 3’ |
| AN89 (F) | Semi-nested PCR | 5’ CCA GCA CTG ACA GCA GYN GAR AYN GG 3’ |
| AN88 (R) | Semi-nested PCR | 5’ TAC TGG ACC ACC TGG NGG NAY RWA CAT 3’ |

Supplementary Methods:

*Dynamic range determination for E11*

Before calibrating the ICC-NanoporeSeq method, an experiment was performed to establish the linear range of amplification for a selected EV type, E11, which amplifies well in RD cells. For these experiments, 6-well plates of fully confluent RD cells were infected with serial dilutions of an E11 suspension in maintenance media in triplicate wells. Immediately after inoculation, one well was scraped, and its entire contents (cells and virus suspension) were frozen at -20°C (t0). The other two wells were incubated for 24 hours at 37°C before being scraped and frozen at -20°C (t24). The samples were centrifuged for 10 minutes at 1000 *g* to remove cell debris, the supernatant was transferred into a clean tube and RNA was extracted from 140 μL of supernatant as described in the main text. After RNA extraction, the concentration of E11 was quantified at both t0 and t24 using the pan-enterovirus RTdPCR assay described in the main text, and results were used to determine the linear range of amplification.

These experiments showed that E11 could be efficiently amplified in RD cells and quantified by RTdPCR over an input range of 7x10¹ to 7x10⁵ MPN/ml (Supplementary Figure S1), consistent with the dynamic range previously reported for ICC-RTqPCR^2^. Within this range, genome copy numbers increased proportionally with input concentration, whereas a plateau was observed at higher concentrations, indicating saturation of amplification.

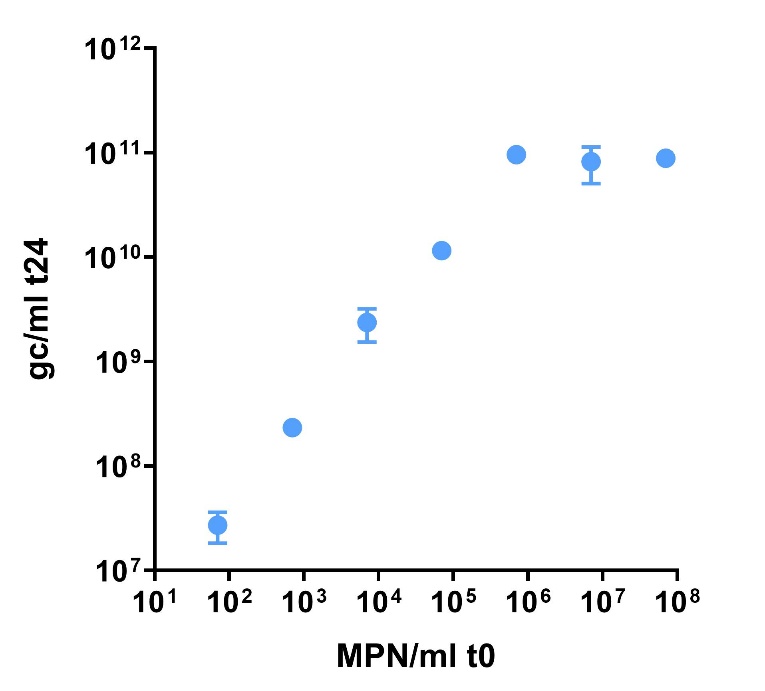

**Supplementary Figure S1.** Dynamic range of amplification of E11 after 24 hours of amplification on RD cells, measured by pan-enterovirus RTdPCR.

*Assessment of type-specific replication in RD cells*

To assess type-specific replication in RD cells, a mixture containing equal proportions of the ten EV types was prepared based on infectious concentration (MPN/ml) and used to inoculate confluent RD cells. The mixture was tested at input concentrations of 10², 10³ and 10⁴ MPN/ml. For each concentration, one well was scraped immediately after inoculation, while two additional wells were incubated for 20 hours at 37°C before being scraped. For the 10^4^ MPN/ml concentration, the experiment was performed in triplicate. Samples were frozen at -20°C and, following centrifugation, RNA was extracted from each sample. VP1 amplicons were subsequently generated and Nanopore sequencing was performed as described in the main text. For each EV type, Nanopore read counts were normalized to the number of reads obtained for the E3 spike-in control. The change in abundance over the course of 20 hours was calculated as the log fold-change of E3-normalized read counts, with each incubated replicate compared independently with the single initial sample obtained at the corresponding input concentration. EV types showing a log fold-change > 0, corresponding to an increase in normalized read counts over the course of 20 hours, were considered to show evidence of replication in RD cells (Supplementary Figure S2). Only EV types meeting this criterion were subsequently included in the generation of type-specific calibration curves (CVB5, CVA9, E6, E11, E25 and E30).

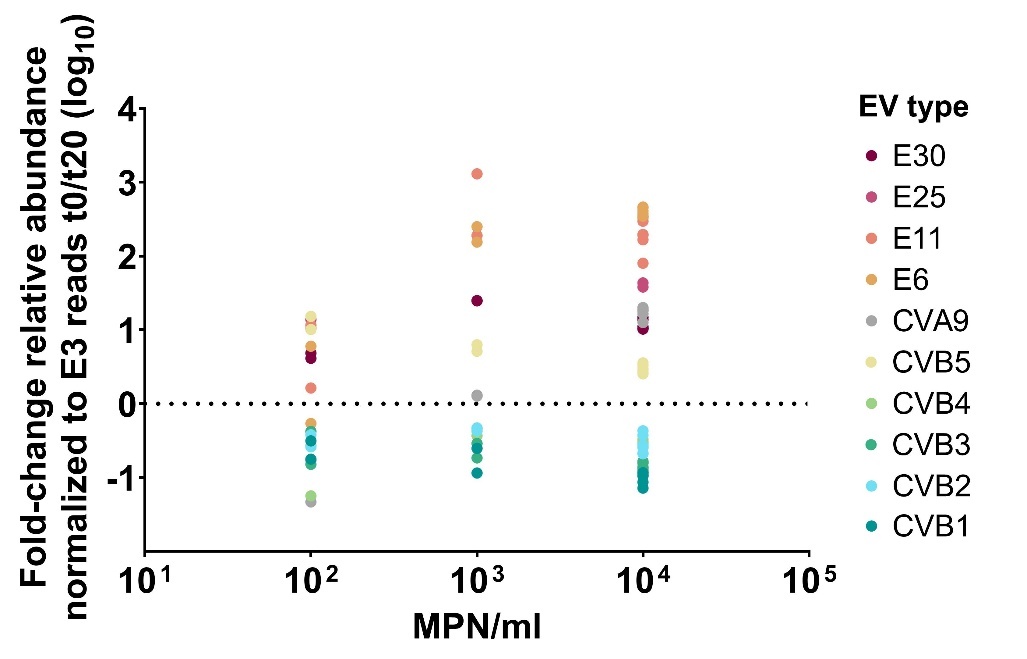

**Supplementary Figure S2.** Assessment of type-specific replication of EVs in RD cells. Scatter plot showing the log fold-change in E3-normalized read counts over the course of 20 hours for each EV type at different input infectious concentrations (MPN/ml). A log fold-change > 0 indicates an increase in E3-normalized read counts over the course of 20 hours and was used as the criterion to identify EV types showing evidence of replication in RD cells.

Sanger sequences for the different EV types used in the study:

>CVB1_AN89

ACTATGCAGACAAGGCACGTGAAGAATTATCATTCCAGATCTGAATCAACCATTGAAAACTTCCTGTGCCGATCCGCTTGTGTTTACTATGCTACCTATACAAATAACACAGAAAAAGGGTATGCAGAGTGGGTCGTAAACACTAGGCAGGTAGCCCAATTAAGAAGAAAGCTGGAGCTGTTCACCTATTTAAGATTTGATTTGGAGCTGACATTTGTGATAACGAGCGCTCAACAACCCAGCACTGCTACTAGTGTGGACGCCCCTGTGCAAACGCACCAGATTATGTACATCCCCCCAGGT

>CVB2_AN89

AGACACGTGCATAACTATCATTCAAGGTCAGAGTCCAGCGTGGAAAATTTCCTGGCGCGATCGGCGTGTGTGTTCTATACAACGTACACAAATAGTAAAAACGCCGCAAAGGAGAAAAAGTTTGCAACATGGAAAGTCAGTGTTAGGCAGGCCGCACAATTAAGAAGGAAGTTGGAACTGTTCACATACTTGCGCTGTGACATCGAGCTCACATTTGTTATCACCAGCGCACAAGATCCATCGACTGCAACCAATTTGGATGTACCAGTGTTGACCCACCAAATAATGTACATCCCCCCAGG

>CVB3_AN89

TGCAAACGCGTCATGTGAAAAATTACCACTCGAGGTCCGAGTCAACAATTGAGAACTTTGTATGCAGATCTGCATGTGTTTATTTCACAGAGTACGAAAATTCAGGGTCAAATCGGTATGCTGAATGGGTGATAACAACTCGCCAGGCAGTGCAATTGAGAAGAAAGTTAGAGTTCTTCACATACATGAGGTTTGATCTAGAATTAACCTTTGTCATCACGAGTACCCAGCAACCCTCCACAACCCAGAACCAAGATGCTCAGATCCTCACCCACCAAATAATGTACATCCCCCCAGGTG

>CVB4_AN89

TGCAAACCAGACACGTGCACAACTACCACTCAAGGTCAGAATCGTCAATTGAGAACTTCTTGTGTAGATCTGCATGTGTAATTTACATTAAATATTCAAGCGCTGAGTCTAACAACTTAAAGCGTTATGCAGAGTGGGTCATTAATACAAGACAGGTGGCACAGCTGCGCCGGAAAATGGAAATGTTCACATACATTCGCTGCGACATGGAATTGACATTTGTCATAACCAGCCATCAGGAAATGTCTACAGCTACCAACTCCGATGTTCCAGTTCAAACACACCAAATAATGTACATCC

>CVB5_AN89

CTATGCAAACCAGACATGTGAAAAACTACCACTCGAGATCGGAGTCGACAGTGGAAAACTTTCTGTGTAGATCAGCATGCGTGTTTTACACCACATACAAAAATCATGGTACTGATGGTGACAACTTTGGTTACTGGGTGATCAACACACGCCAGGTGGCTCAACTACGGCGTAAGCTTGAGATGTTCACATATGCGAGATTTGATCTCGAGCTGACCTTTGTGATCACGAGCACTCAAGAACAATCCACCATACAAGGCCAAGATTCACCAGTGCTCACCCATCAAATCATGTACATCCCCCCA

>CVA9_AN89

GACACCATGCAGACTAGGCACGTGAAGAATTACCATACTCGGTCTGAGTCCACCGTGGAAAACTTTCTTGGCAGATCAGCCTGTGTTTATATGGAGGAATACAAGACCACAGATAATGATGTCAACAAGAAATTCGTGGCGTGGCCAATCAACACTAAACAGATGGTGCAAATGCGTAGGAAGCTAGAGATGTTCACCTACCTCAGATTTGACATGGAAGTAACCTTTGTGATCACAAGTCGGCAAGATCCTGGAACCACACTAGCACAAGACATGCCAGTGTTGACGCACCAGATTATGTACATCCCCCCAGGTGGGTCCAGTAA

>E6_AN89

CATACAAACGCGCCACGTCAAAAATTTTCACGTGAGGTCAGAGTCATCAGTGGAAAACTTTCTCAGTAGGTCTGCCTGTGTGTACATCGTGGAGTATAAGACAAGAGATAACACTCCAGACAAGATGTACGACAGCTGGGTCATCAACACCAGACAAGTTGCCCAGTTGCGTAGGAAATTGGAATTCTTCACCTATGTCAGGTTTGATGTGGAAGTCACGTTTGTTATCACCAGTGTGCAAGATGACTCAACCAGACAAAACACCGATACACCGGCTCTCACACACCAGATAATGTACATTCCCCCAGGTGGGTCCAGTAA

>E11_AN89

ATGCAACCAGGCATGTCAAGAACTACCATTCCAGATCTGAGTCCAGCATTGAAAACTTCCTCAGCAGATCTGCCTGCGTTTATATGGGAGGATACCACACAACCAACACTGACCAGACAAAATTATTTGCCTCATGGACTATTAGTGCACGACGCATGGTTCAAATGAGACGCAAGCTAGAGATCTTCACTTACGTCCGTTTTGATGTGGAGGTGACTTTTGTGATTACCAGCAAGCAGGACCCGGGCAACCGATTGGGCCAAGACATGCCACCCCTGACTCACCAGATCATGTACATCCCCCCAGGT

>E25_AN89

AATGCAAACCAGACATGTTGTTAACCACCACATTAGGTCAGAATCCTCAATAGAGAACTTTTTAAGTAGATCAGCGTGTGTCTACATTGACGTGTATGGTACAAAGGAGAATGGTAACATTGAACGCTTTACCAACTGGAAGATCAACACACGTCAAGTTGTTCAGTTAAGGCGCAAGCTGGAAATGTTTACGTACATCAGATTTGATGTGGAGATTACATTCGTGATCACAAGCACCCAGGGGACATCAACCCAAACGAACACAGACACCCCCGTGCTCACACATCAGGTAATGTACATTCCCCCAGGTGGGTCCAGTAA

>E30_B1_AN89

ACAATGCAGACACGGCACGTGGTCAACTACCATACCAGATCAGAATCGTCAATAGAGAACTTTATGGGTAGAGCAGCGTGTGTGTACATCGCCCAGTACGCCACAGAGAAGGTCAATGACGAGTTAGACAGGTACACCAACTGGGAAATAACAACCAGGCAAGTGGCACAATTGAGACGGAAACTGGAAATGTTCACATACATGAGATTTGACCTTGAGATCACATTTGTCATCACCAGTTCCCAGCGCACTTCAACCACATATGCATCGGATTCCCCTCCACTAACGCACCAAGTGATGTACATCCCCCCAGGTGG

>E30_B2_AN89

CATGCAGACACGGCACGTGGTCAACTACCATACCAGATCAGAATCGTCAATAGAGAACTTTATGGGTAGAGCAGCGTGTGTGTACATCGCCCAGTACGCCACAGAGAAGGTCAATGACGAGTTAGACAGGTACACCAACTGGGAAATAACAACCAGGCAAGTGGCACAATTGAGACGGAAACTGGAAATGTTCACATACATGAGATTTGACCTTGAGATCACATTTGTCATCACCAGTTCCCAGCGCACTTCAACCACATATGCATCGGATTCCCCTCCACTAACGCACCAAGTGATGTACATCCCCCCAGGTGGTCCAGTAA

>E3_AN89

ATGCAACGAGGCATGTCAAGAATTACCACTCTAGGACTGAATCATCCATAGAGAACTTCTTGTGTAGAGCGGCATGTGTTTACATCACAACCTATAAGTCGGCTGGTGGTACCCCCACGGAACGATACGCTAGCTGGAGGATCAACACCCGGCAAATGGTGCAGCTGAGAAGAAAATTCGAGCTGTTTACATACTTACGGTTCGATATGGAAATAACATTTGTAATCACCAGTACACAAGACCCAGGGACGCAGCTGGCACAAGATATGCCTGTGTTGACCCATCAGATTATGTACATCCCCCCA
